# Effects of background food on alternative grain uptake and zinc phosphide efficacy in wild house mice

**DOI:** 10.1101/2021.10.11.463604

**Authors:** Steve Henry, Peter R. Brown, Nikki Van de Weyer, Freya Robinson, Lyn A. Hinds

## Abstract

**BACKGROUND:** House mice (*Mus musculus*) cause significant, ongoing losses to grain crops in Australia, particularly during mouse plagues. Zinc phosphide (ZnP) coated grain is used for control, but with variable success. In a laboratory setting, we tested if mice would (1) switch from consumption of one grain type to another when presented with an alternative, and (2) consume ZnP-treated grains when presented as a choice with a different grain.

**RESULTS:** Mice readily switched from their background grain to an alternative grain, preferring cereals (wheat or barley) over lentils. Mice readily consumed ZnP-coated barley grains. Their mortality rate was significantly higher (86%, n=30) in the presence of a less-favoured grain (lentils) compared to their mortality rate (47%, n=29; and 53%, n=30) in the presence of a more-favoured grain (wheat and barley, respectively). Mice died between 4-112 h (median = 18 h) after consuming one or more toxic grains. Independent analysis of ZnP-coated grains showed variable toxin loading indicating that consumption of a single grain would not guarantee intake of a lethal dose. There was also a strong and rapid behavioural aversion if mice did not consume a lethal dose on the first night.

**CONCLUSIONS:** The registered dose rate of 25 g ZnP/kg wheat; ∼ 1 mg ZnP/grain in Australia needs to be re-evaluated to determine what factors may be contributing to variation in efficacy. Further field research is also required to understand the complex association between ZnP dose, and quantity and quality of background food on efficacy of ZnP baits.

## 1 INTRODUCTION

Wild house mice (*Mus musculus domesticus*) cause ongoing and significant problems in grain growing areas of Australia [1]. The primary impact of mice is through damage to crops leading to major economic losses, with damage being particularly acute during mouse plague events that can occur every three to five years with resultant losses exceeding $100 million [1,2,3,4,]. Mice also cause serious damage to intensive livestock and horticulture industries, significant social and environmental damage throughout rural communities through damage to houses, infrastructure and equipment [2], and are a source of zoonotic diseases [5]. Effective management is required to minimise the economic impact caused by mice.

In Australia, acute rodenticides, such as strychnine and zinc phosphide (Zn_3_P_2_; herein ZnP), have been used extensively to manage mouse problems in grain-growing regions [6]. In 1996, ZnP was proposed for use in Australia to replace strychnine as a broad-acre rodenticide for control of mouse populations [7] and to reduce non-target impacts. In May 2000, ZnP-coated wheat grain became the only registered in-crop rodenticide bait (APVMA, https://apvma.gov.au/node/14101) for the management of over abundant mice in broad scale crops in Australia [6,8,9]. For the registered rate of 25 g ZnP/kg wheat bait there is approximately 1 mg ZnP per grain of wheat, meaning theoretically that a single grain is equal to one lethal dose for a 20 g mouse [10]. ZnP bait is applied at 1 kg per hectare resulting in approximately 2-3 grains of bait per square metre. Zinc phosphide is commonly used to control rodent pests in many countries in a range of crops, not just broadacre grain crops, including sugarcane [11], alfalfa [12], coconut [13,14], rice [14,15,16], and fields and forests [17,18]. The applicability and use of ZnP baits across a range of crop types is well illustrated.

The effectiveness of ZnP baits has been highly variable, however, with field studies reporting between 50% to 95% efficacy [8,9,19,10]. When mouse abundance is high and mouse damage is likely, particularly at sowing of crops, growers will often contravene label rates (applying baits at rates > 1 kg/ha) or repeatedly apply baits at short intervals (Henry S and Brown PR, unpublished). This bait application misuse might also contribute to poor efficacy because of behavioural avoidance from sub-lethal doses. Other reasons for variability in efficacy could be related to the bait substrate used and the quantity and/or quality of alternative food available.

Significant changes in farming systems over the past 20 years may also be contributing to the reported changes in the efficacy of ZnP. Before 2002, conventional cropping systems followed a three-to four-year crop rotation, this involved numerous passes with ploughing equipment to control weeds, seed bed preparation and leaving fields fallow in the year prior to planting the next crop. The adoption of conservation agriculture (CA) systems (also known as zero or no-till cropping systems) involves retaining stubble, using herbicides to control weeds and practicing single-pass sowing with disc or narrow tyne to minimise soil disturbance [21]. This has created an environment that is more favourable for mice than conventional cropping systems [22]. In contrast to conventional cropping systems, the habitat in CA systems is more complex with crops growing in amongst crop residue and standing stubble, and in some years high levels of residual grain from the previous crop remaining on the ground. This complexity also means that the ability of mice to detect ZnP bait spread at 1 kg/ha (approximately 3 grains/m^2^) may be reduced when compared to conventional systems where all crop residues are buried by tillage and bait is spread onto bare soil.

Zinc phosphide is currently coated onto wheat grains. Frequent anecdotal reports of poor effectiveness of ZnP make it worth exploring whether an alternative to wheat as a bait substrate should be considered to improve palatability and/or efficacy, especially given complex CA cropping systems with a wide range of crop types (Henry S and Brown PR, unpublished). There have been many studies directed at feeding preferences of house mice, because it is critical to have a bait that rodents will eat. Robards and Saunders [23] reported that house mice preferred soft wheat varieties, canary seed (*Phalaris canariensis*) and rice (*Oryza sativa*), noting however, that canary seeds are relatively small and would be impractical for use in field conditions. Rowe et al. [24] conducted food preference trials for the development of poison baits for mice and found that canary seed, pinhead oatmeal and wheat were well accepted by mice. Pennycuik and Cowan [25] used a small maze to determine odour preferences and showed that mice preferred canary seed or maize rather than a control diet of mouse breeder pellets. A further study examining flavour preferences among wheat varieties indicated mice had preferences for hard white spring wheat varieties over hard red spring wheat varieties [26]. Other recent work has successfully investigated volatile food attractants to improve baits for trapping rodents (e.g.: [27]). Overall, the choice of food by mice appears to be dependent on its palatability and particle size.

The objective of this study was to identify an alternative grain type to wheat that may be more attractive, readily detected and palatable to mice in the presence of background food, as might be found in complex habitats associated with CA cropping systems. Commercial ZnP is currently mixed on wheat grains, but could the use of alternative grains, such as high protein, hard wheat (durum), malt barley or lentils, be more attractive to mice? Attractiveness might be important especially when bait is applied in the presence of different background food types (i.e. previous crop type). We addressed two questions in this laboratory study: (1) Do mice switch from consumption of a background grain type when presented with a choice of an alternative grain type? (2) Do mice consume the alternative grain type when it is coated with ZnP and presented as a choice with different background grains? The identification of alternative, palatable grain bait substrates could provide growers and the grains industry with a selection of substitutes to wheat and could improve management of house mice in crops.

## 2 METHODS

### 2.1 Animals

Wild house mice were trapped in cropping paddocks near Walpeup in the Central Mallee of Victoria (35°06’S, 142°01’E). Single capture Longworth traps (Longworth, Abingdon, UK) were set adjacent to a wheat stubble paddock and trapped mice were weighed, sexed and checked for alertness and general body condition. Only healthy adult mice (n = 90) were transported to an animal holding facility at CSIRO in Canberra, ACT, Australia. Mice were individually housed in mouse cages (L × W × H, 26 × 40 × 20 cm) containing wood shavings for bedding, tissue for nesting material, a cardboard tube for shelter, and *ad libitum* food and water. Mouse cages were placed on racks in a temperature-controlled room (22°C ± 3°C) and lights were set at 12 h Light: 12 h Dark (On at 0600 h and off at 1800 h each day). Mice were acclimatised to facility conditions for two weeks while on a maintenance diet of standard laboratory mouse pellets (Gordon’s Specialty Stock Feeds, NSW) before and between Experiments 1 and 2. The trapping and use of animals in experiments was approved by the CSIRO Wildlife and Large Animal - Animal Ethics Committee, Approval No. 2018-22.

### 2.2 Experimental design

The same mice were used for the two experiments. We define ‘background’ food as the maintenance food provided to mice (as a reflection of naturally available food sources available in fields), ‘alternative’ food or grain as a food substance which is provided to mice as an alternative challenge food type to the background food, and ‘substrate’ in reference to the grain type that carries toxic grains. The aim of Experiment 1 was to identify food preferences of house mice given an alternative grain choice in the presence of different background grain types. The results would establish potential alternative bait substrate candidates for use in Experiment 2 which aimed to determine the willingness of mice to consume different toxic ZnP-coated grains in the presence of a single alternative grain type.

### 2.3 Experiment 1: Do mice switch their consumption of a background grain type when presented with the choice of an alternative grain type?

Following the initial acclimation period, mice were randomly allocated by weight and sex into three treatment groups, (n = 30 mice per group, Table 1) and their diet of laboratory pellets was replaced with a background grain, either common wheat, barley or lentils *ad libitum*, for two weeks (Table 2). These three grain types putatively may be more attractive due to a higher protein or sugar content (Table 2) and are able to be distributed using a standard bait spreader. The grains are also representative of those that mice commonly encounter in the Victorian and South Australian cropping regions. After two weeks on background grain, each group of 30 mice were further sub-divided into three groups (n = 10 mice per group, 5 males, 5 females), balanced by body weight (12.8-18.0 g females; 11.5-20.0 g males), to establish nine groups of mice.

**Table 1.** Treatment groups (n = 9) received one of three background grain types for two weeks and then a choice of one alternative grain types for five days for Experiment 1. Each treatment group comprised 10 mice (5 males, 5 females).

| Treatment | Mice | Background food type | Alternative food type |
| --- | --- | --- | --- |
| 1 | 10 (5 ♂, 5 ♀) | Common wheat | Durum wheat (high protein) |
| 2 | 10 (5 ♂, 5 ♀) | Common wheat | Malt Barley with husk |
| 3 | 10 (5 ♂, 5 ♀) | Common wheat | Lentils |
| 4 | 10 (5 ♂, 5 ♀) | Barley with husk | Durum wheat (high protein) |
| 5 | 10 (5 ♂, 5 ♀) | Barley with husk | Malt Barley with husk |
| 6 | 10 (5 ♂, 5 ♀) | Barley with husk | Lentils |
| 7 | 10 (5 ♂, 5 ♀) | Lentils | Durum wheat (high protein) |
| 8 | 10 (5 ♂, 5 ♀) | Lentils | Malt Barley with husk |
| 9 | 10 (5 ♂, 5 ♀) | Lentils | Lentils |

**Table 2.** Protein, sugar, energy (kJ), dietary fibre and total carbohydrate content (g/100 g) for each grain type (USDA, https://fdc.nal.usda.gov/fdc-app.html#/food-search, FSANZ, https://www.foodstandards.gov.au/science/monitoringnutrients/afcd/Pages/fooddetails.aspx).

| Grain type | Protein | Sugar | Energy (kJ) | Dietary fibre | Total carbohydrate |
| --- | --- | --- | --- | --- | --- |
| Common wheat, *Triticum aestivum* | 11.5-15.1 | 0.4-1.3 | 1367 | 10.6-11.4 | 58.3-71.2 |
| Durum wheat, *Triticum durum* | 13.7 | 0.5 | 1420 | - | 71.1 |
| Barley, *Hordeum vulgare* | 10.1-12.5 | 0.8-1.0 | 1428-1480 | 13.1-17.3 | 60.6-73.5 |
| Barley Malt | 10.3 | 0.8 | 1510 | 7.1 | 78.3 |
| Lentil, *Lens culinaris* | 23.0-23.9 | 1.8-2.7 | 1364-1500 | 10.8-13.7 | 45.7-63.1 |

For the next five days the nine groups of mice were provided with a choice of their background grain and an alternative grain type, either durum wheat, malt barley with husk or lentils (Table 1). The amount of each grain offered equated to approximately 10% by body weight. The different grains were dyed either red or green using a tasteless, odourless food dye (Queen Fine Foods, food colouring) to facilitate sorting of remaining grains each day to determine amount consumed.

During the 5-day experimental period each mouse cage contained only a cardboard shelter tube and two food dishes, one on each side of the cage and secured to the cage floor using “Blu Tack” (Bostik Australia Pty Ltd, Thomastown, Victoria, Australia) to minimise spillage. Each morning between 0800 h and 1030 h mice were removed from their cage and weighed. Cages were cleaned and remaining grain was sorted based on colour and dried in a drying oven at 40°C for 24 h. At 1600 h food dishes containing a known amount of each grain (at least 10% body weight) were added. The position of each grain type was reversed daily to prevent mice habituating to a particular food dish.

Control food dishes (n = 6) holding each grain type were set up in an empty cage with attached water bottle and placed on racks adjacent to trial animals. The following morning control samples were dried at 40°C for 24 h. Changes in control grain weight (gain or loss) were used as a correction factor when calculating the amount of grain consumed by trial mice. The amount of each grain type eaten each day was calculated by subtracting the amount remaining (weight corrected after drying for 24 h), from the original amount of food provided.

### 2.4 Experiment 2: Do mice consume the alternative grain type when it is coated with ZnP and presented as a choice with different background grains?

At the conclusion of Experiment 1, all mice were returned to a diet of standard laboratory mouse pellets for two weeks. The original three groups of 30 mice were then allocated to a different background grain type for a further two weeks prior to the commencement of Experiment 2. Each mouse cage was set-up with a shelter tube and two secured food dishes as described previously for Experiment 1.

Experiment 2 was conducted over six days. For three days, at 1600 h, animals were provided with their background grain type (at least 10% body weight) in one food dish, and ZnP-coated grains (n = 10) in the other food dish (Table 3). As in Experiment 1, the location of the food types in the cage was reversed each day. ZnP-coated grains were prepared independently according to the registered commercial application rate (25 g ZnP/kg).

**Table 3.** Treatment groups (n = 9) received one of three background grain types for two weeks and then a choice of one alternative grain types for five days for Experiment 2. Each treatment group comprised 10 mice (5 males, 5 females). Mice were provided with a different background food type from Experiment 1.

| Treatment | Mice | Non-toxic treatment<br>Background food type | Toxic treatment<br>Zinc phosphide treatment |
| --- | --- | --- | --- |
| 1 | 10 (5 ♂, 5 ♀) | Wheat | Barley with husk |
| 2 | 10 (5 ♂, 5 ♀) | Wheat | Malt barley with husk |
| 3 | 10 (5 ♂, 5 ♀) | Wheat | Malt barley without husk |
| 4 | 10 (5 ♂, 5 ♀) | Barley with husk | Barley with husk |
| 5 | 10 (5 ♂, 5 ♀) | Barley with husk | Malt barley with husk |
| 6 | 10 (5 ♂, 5 ♀) | Barley with husk | Malt barley without husk |
| 7 | 10 (5 ♂, 5 ♀) | Lentils | Barley with husk |
| 8 | 10 (5 ♂, 5 ♀) | Lentils | Malt barley with husk |
| 9 | 10 (5 ♂, 5 ♀) | Lentils | Malt barley without husk |

The results for Experiment 1, indicated mice had a slight but not significant preference for malt barley over lentils and durum wheat. However, during Experiment 1 mice offered a grain in a husk (barley or malt barley) were observed removing the husk prior to consumption of the grain. This raised concerns about using husked grain as an alternative bait substrate, as the ZnP mixture is coated on the outside of the grain and by de-husking the grain the mice may not consume a lethal dose of ZnP. Therefore, barley with the husk, malt barley with the husk, and malt barley without the husk were coated with ZnP for testing in Experiment 2 to determine if the husk made a difference to the effectiveness of the bait (Table 3).

Following addition of the toxic and non-toxic grains, mice were monitored every 30 minutes for six hours (1600-2200 h). The number of ZnP-coated grains consumed, and the activity and condition of each mouse was recorded. Mice showing clinical signs of ZnP poisoning (moribund, loss of hindquarter function, lateral or sternal recumbency) were humanely killed using an overdose of isoflurane (Laser Animal Health, Pharmachem, Eagle Farm Queensland). The time to death (humane killing) from addition of toxic grains was recorded. For mice that did not display signs of ZnP poisoning by 2200 h, hourly monitoring re-commenced the following morning from 0700 h. At 1600 h each day, any remaining ZnP grains (whole or partially consumed) were counted and replaced with 10 new toxic ZnP grains. Remaining background food was removed and dried at 40°C for 24 h before being weighed. Control grains (n = 6 replicates) were set up as in Experiment 1 and used as a correction factor for moisture loss of grains. Any animal still alive three days after initial presentation with toxic ZnP grains was checked twice daily for a further three days, then humanely killed.

### 2.5 Analysis of ZnP on grains

A sample of ZnP-coated grains (n = 10) and uncoated grains (n = 10) of each grain type were analysed by Agrifood Technology, Werribee, Victoria. Individual grains were weighed into a digestion tube and 5 ml nitric acid added. Each tube was sealed and placed in the MARS microwave digestion unit and digested at 170°C under pressure. Once cooled the sample was diluted to 20 ml, filtered and analysed using the Inductively Coupled Plasma Excitation (AS 3641.2 – 1999) method. A standard calibration curve was prepared using six standards. Blank samples were included as well as a spiked reference wheat sample to determine recoveries of Phosphorous (P) (99%) and Zinc (Zn) (101%). Continuous Calibration Verification and Quality Controls were included after every 10 samples and at the end of the sequence and were found to be within specifications (± 10%). Any samples greater than the calibration range were diluted to within the range and re-run.

### 2.6 Statistical Analysis

For Experiment 1, for each grain type the amount consumed was calculated as the difference between the amount provided and that remaining after correcting for changes in moisture. The proportion of each alternative grain and background grain consumed by individual mice each day (days one to five) was calculated. A value of “0” indicates strong preference to background grain, “0.5” indicates no preference and “1” indicates strong preference for alternative grain. Some values were negative due to potential moisture correction error, to deal with this any negative values were adjusted to equal zero. All data were then transformed ((*Y* × (*length*(*Y*) – 1) + 0.5) / *length*(*Y*), where *Y* is the proportion of alternative grain taken) to account for real zero values prior to a logit transformation.

A linear mixed effect model in R (R version 1.1.456, [28]) using the *lmer* function in the lme4 package [29] was used to examine differences in the logit transformed proportion of alternative grain take as the dependent variable between treatment groups. The data were modelled with each combination of background grain type and day as fixed effect factors, and individual animals nested in treatments as random effect factors. Confidence intervals (CI) were extracted from the model using the *confint* function and back transformed. The overlap in confidence intervals at a 95% confidence level were used to evaluate effect size of alternative grain consumption for each treatment group. Confidence intervals (CI) are reported as 95% CI [LL, UL], where LL is the lower limit of the CI and UL is the upper limit. Values reported in figures are extracted from the model using the *fixef* function and back transformed to real values.

For Experiment 2, linear regression models in R (R Core Team 2020) were used to compare the number of toxic grains consumed for each combination of toxic grain type, background grain type, night, and percent mortality with individual animals as random effect factors. Reported *F*-values and *P*-values were obtained by an analysis of variance (ANOVA) of the fitted linear regression models. Means are reported ± 1 standard deviation (SD) throughout.

## 3 RESULTS

### 3.1 Experiment 1: Do mice switch their consumption of a background grain type when presented with choice of an alternative grain type?

When mice were provided with the different background grain types (lentils, barley with husk or wheat) they maintained their body weight (Females: 14.8 ± 1.2 g; Males: 16.2 ± 2.2 g) and general body condition.

When provided with the choice of an alternative grain type (durum, malt barley with husk or lentils), the background grain type strongly influenced the proportion of alternative grain consumed by mice (Figure 1). There was no difference in the proportion of the alternative grain consumed between days one to five for any treatment group (*F*_*4, 404*_ = 1.44, *P* = 0.22).

**Figure 1.**
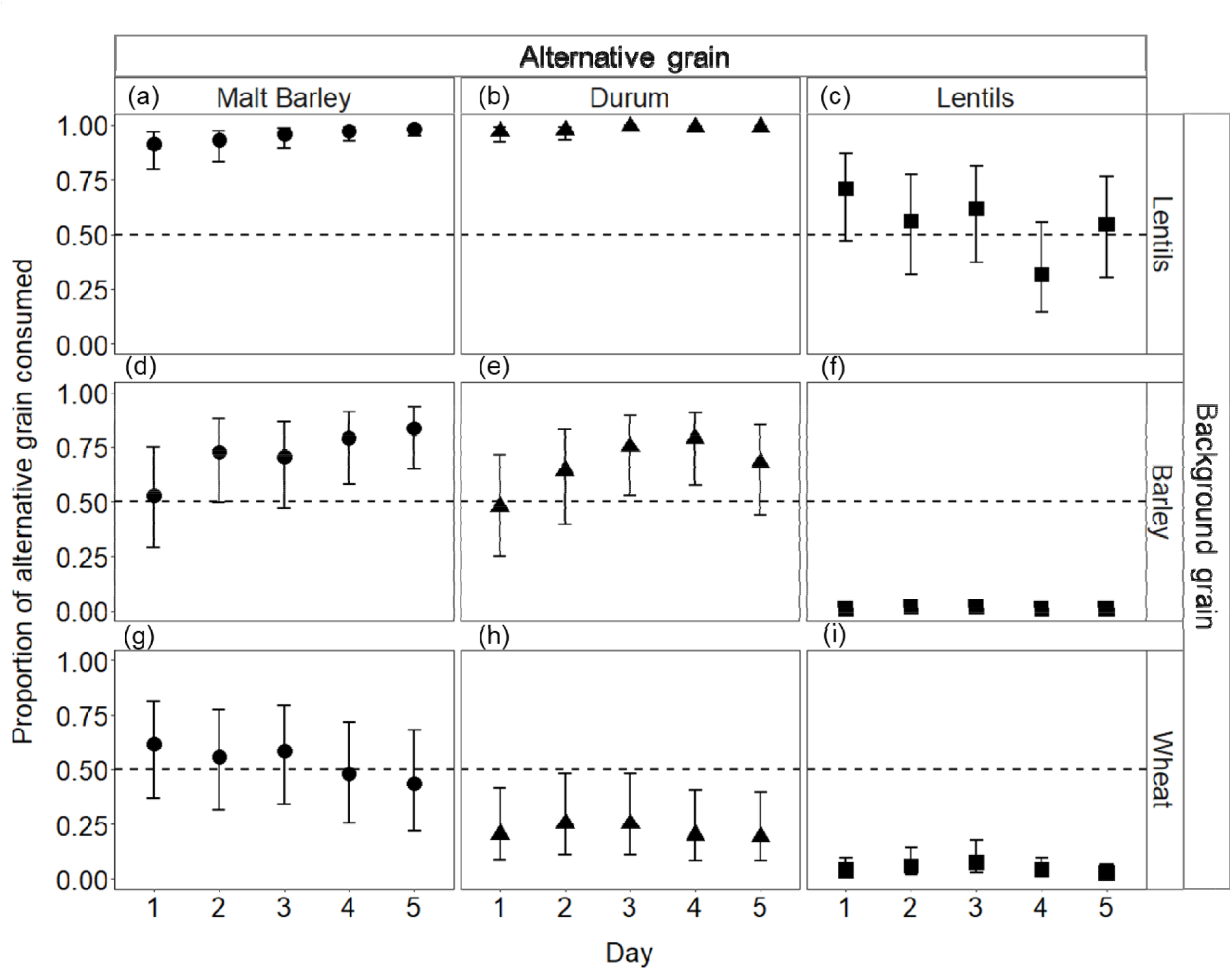
The proportion of alternative grain (a, d, g - malt barley with husk (⍰); b, e, h - durum wheat (▴); c, f, i - lentils (▪)) consumed by mice (n = 10 per group) established on different background grains (a, b, c - lentils; d, e, f – barley with husk; g, h, i - common wheat) over 5 days. A value of “0” indicates strong preference to background grain, “0.5” indicates no preference (represented by dashed line) and “1” indicates strong preference for alternative grain. Shapes (⍰, ▴, and ▪) represent estimates of fixed effects (individual mice as random effects), error bars represent 95% CI.

Mice established on a lentil background and then offered an alternative grain of malt barley with husk (Figure 1a) or durum wheat (Figure 1b) switched to the alternative grain on night one (malt barley mean proportion = 0.91, 95% CI [0.80, 0.97]; durum wheat mean = 0.97, 95% CI [0.93, 0.99]) and maintained that switch for the next four nights. Mice offered lentils as their alternative grain showed no preference (mean = 0.71, 95% CI [0.47, 0.87]) over the five treatment nights (Figure 1c), indicating that the position of the food dishes in the cage did not influence the choice of grain consumed.

Mice established on a barley with husk background tended to display an increasing preference towards alternative grain of malt barley with husk (Figure 1d) and durum wheat (Figure 1e), although they did not completely switch to the alternative malt barley (mean = 0.53, 95% CI [0.29, 0.75]), or durum wheat (mean = 0.48, 95% CI [0.25, 0.71]). When offered lentils as the alternative grain (Figure 1f), a strong preference towards the background barley with husk was observed (mean = 0.009, 95% CI [0.003, 0.02]).

No preference was observed for mice established on a wheat background. When offered alternative malt barley with husk, mice consumed similar proportions of both grain types (mean = 0.62, 95% CI [0.37, 0.81]; Figure 1g). Mice offered durum wheat (Figure 1h) or lentils (Figure 1i) as their alternative grain did not switch and maintained greater consumption of their background wheat (durum wheat mean = 0.21, 95% CI [0.09, 0.41]; mean = 0.04, lentils 95% CI [0.01, 0.10]).

### 3.2 Experiment 2: Do mice consume the alternative grain type when it is coated with ZnP and presented as a choice against different background grains?

During the first night of the trial, all mice consumed background grain. For mice on background lentils, barley with husk and wheat, consumption was 4.9 ± 3.3, 9.7 ± 4.7 and 10.5 ± 4.5 % of their body weight, respectively.

Mice consumed toxic ZnP grains regardless of grain type used. Most mice (n = 87/90) consumed at least one toxic ZnP grain within 30-120 minutes of addition to the cage on the first night. Only one mouse, on a wheat background, did not consume any toxic ZnP grains on any of the three nights they were offered. The number of toxic ZnP grains consumed on the first night by individual mice (n = 87) across all treatment groups was 4.6 ± 3.2 grains (min = 0, max = 10, median = 4 grains). The average number of toxic grains consumed by individual mice (n = 89) over the three-night trial was 4.9 ± 3.2 grains (min = 0, max = 14, median = 4 grains). Consumption of toxic grains by mice was strongly influenced by their background grain type (*F*_*2, 167*_ = 31.6, *P* < 0.001; Figure 2). On the first night of exposure to toxic ZnP grains, mice on a lentil background consumed 7.3 ± 2.5 toxic grains (min = 3, max = 10, median = 6.5 grains), while mice on a barley background consumed fewer toxic grains (4.5 ± 2.9) (min = 1, max = 10, median = 3.5 grains), and those on a wheat background consumed 2.1 ± 1.6 grains (min = 0, max = 7, median = 2 grains). However, there was no difference between the number or type of toxic ZnP grains consumed by mice for each of the background grain type groups (*F*_*4,167*_ = 1.8, *P =* 0.12).

**Figure 2.**
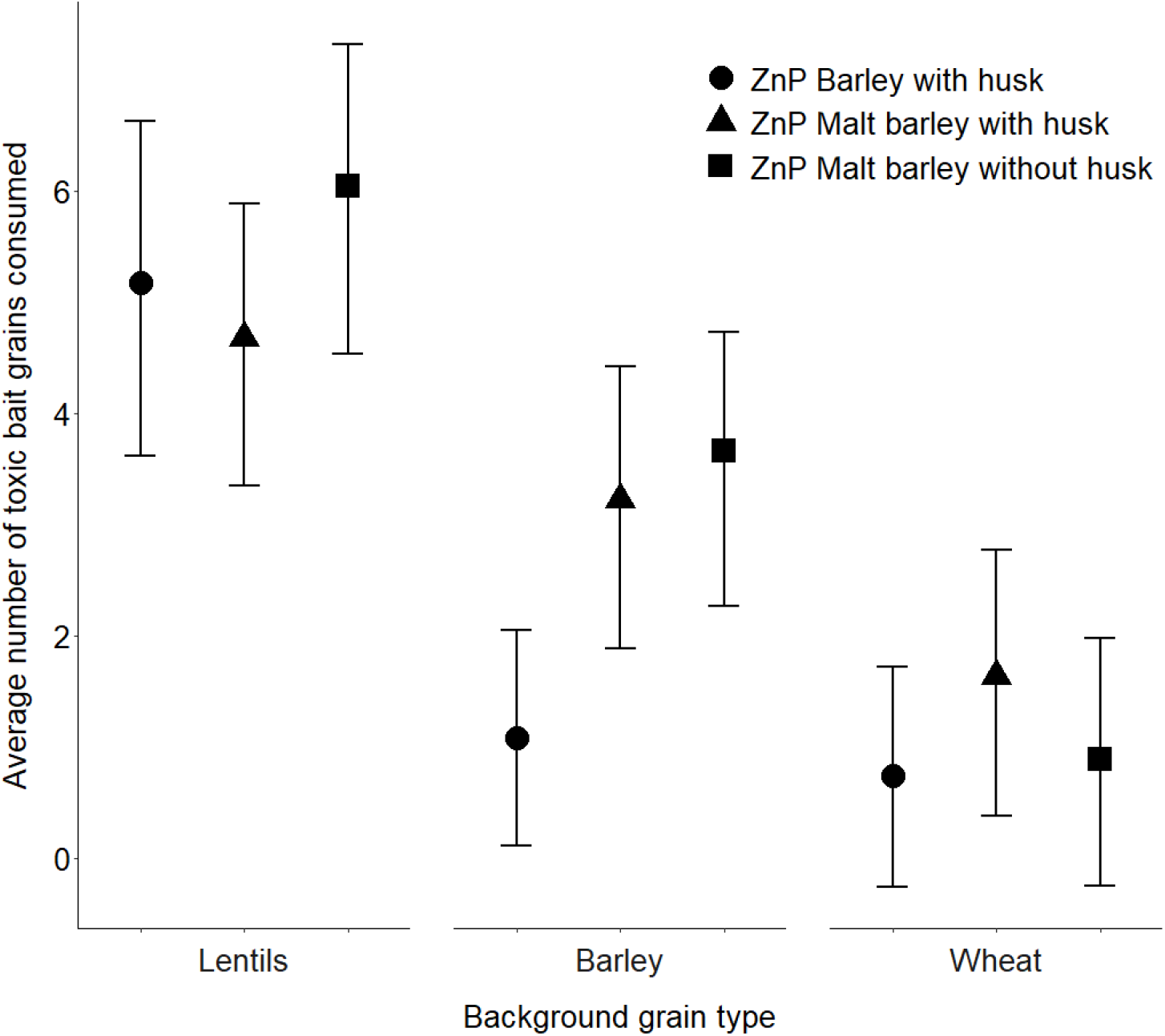
Number (Mean ± 95% CI) of ZnP-coated grains (barley with husk (⍰), malt barley with husk (▴) or malt barley without husk (▪)), consumed by mice (n = 10) over three nights while held on either lentil, barley with husk or wheat background. Shapes (⍰, ▴, and ▪) represent estimates of fixed effects (individual mice as random effects).

Mortality (%) was strongly related to the number of ZnP-coated grains consumed (*F*_*1, 54*_ = 17.6, *P* = 0.0001). Of 10 mice that consumed one ZnP-coated grain, four died (40%). For the 45 mice that consumed between one and four ZnP-coated grains, 18 (40%) died, whereas of the 44 mice that consumed more than four ZnP-coated grains, 38 died (86.4%). The highest mortality (86%) occurred for mice established on a lentil background (Table 4). Three mice, all on a lentil background consumed nine, 10 and 14 ZnP-coated grains but did not die. Across all background grain types, the time to death after consumption of toxic grains for 54 animals ranged between 4 and 66 h (21.2 ± 12.48 h; median = 18 h). Two other animals died at 87 and 112 h, respectively. Overall, 48 animals (85.7%) died within 30 h of exposure to ZnP-coated grains, while eight animals (14.3%) died between 39 and 112 h.

**Table 4.** Mortality (%) and time to death (h) of mice for each background grain type; minimum (Min), maximum (Max), median and mean ± SD time to intervention. n = number of mice humanely killed/total number of mice in treatment group.

| Background grain<br>type | Percent<br>mortality (%) | Time to death (h) |  |  |  |  | n |
| --- | --- | --- | --- | --- | --- | --- | --- |
| | | Min | Max | Median | Mean | ± SD | |
| Lentils | 86.7 | 7 | 47 | 18 | 20.5 | 8.8 | 26/30 |
| Barley | 53.3 | 5 | 66 | 15 | 23.0 | 19.1 | 16/30 |
| Wheat | 48.3 | 4 | 112 | 21 | 31.7 | 30.1 | 14/29 |

For all background grain types, some mice did not die after consuming toxic ZnP grains on the first night of exposure. Each of these mice showed a significant decrease in consumption of toxic ZnP grains on subsequent nights (*F*_*2, 99*_ = 37.8, *P* < 0.001; Figure 3a). Consumption of background grain increased by night three for all surviving mice in all treatment groups (*F*_*2, 99*_ =7.3, *P*= 0.001; Figure 3b).

**Figure 3.**
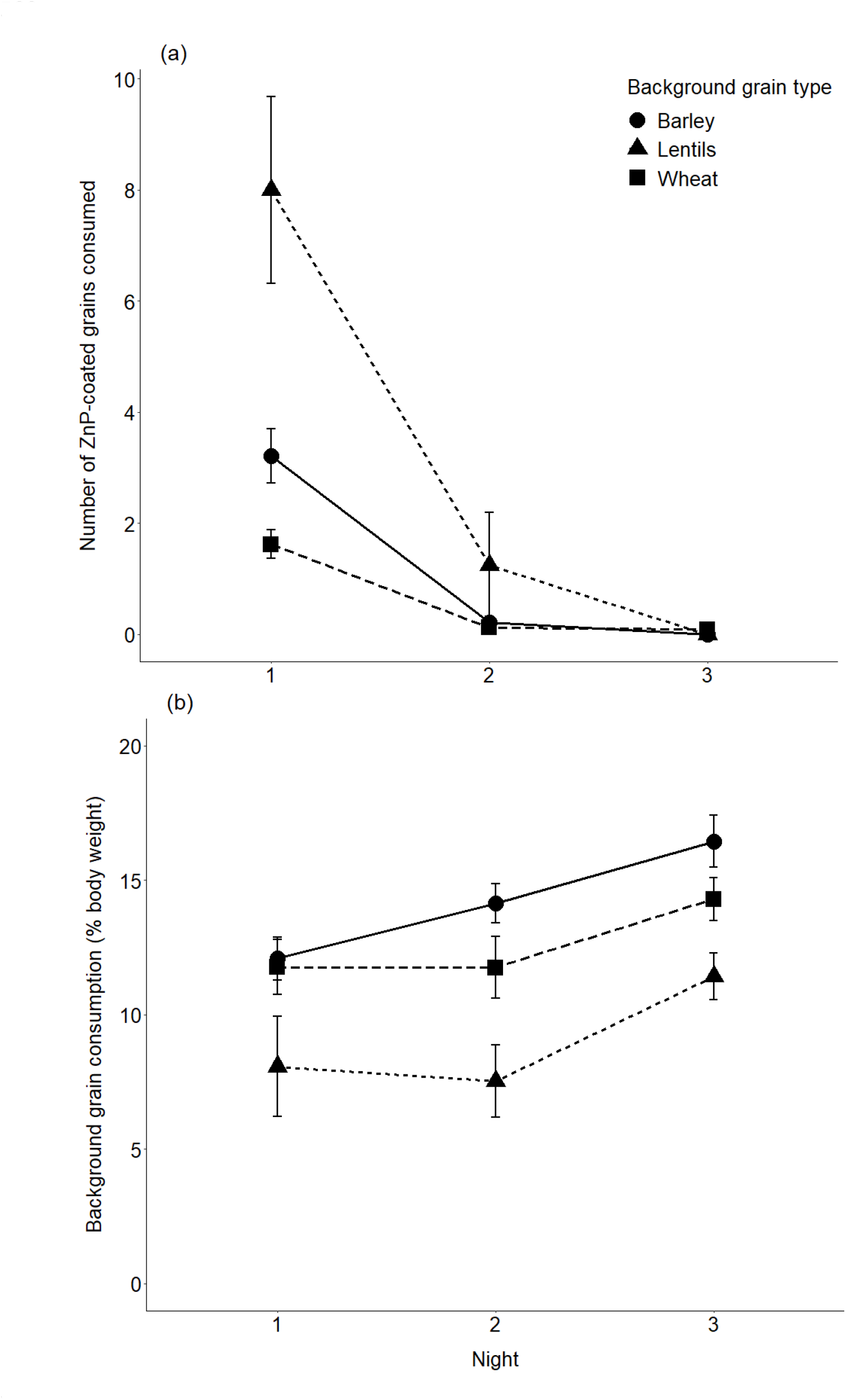
(a) Number (Mean ± SE) of ZnP-coated grains consumed by mice that did not die while held on barley (n = 14 mice), lentils (n = 4 mice) and wheat (n = 16 mice) background food groups over three nights. (b) Amount (% body weight; Mean ± SD) of background food eaten by these surviving mice over three nights.

### 3.3 Toxic ZnP grain analysis

The amount of ZnP on individual grains (n = 30) varied between 0.47 and 1.95 mg ZnP/grain, with an overall average of 0.93 ± 0.4 mg ZnP/grain (Figure 4). For each bait substrate there was an average of 1.26 ± 0.4, 0.78 ± 0.4 and 0.75 ± 0.2 mg ZnP/grain for barley with husk, malt barley without husk and malt barley with husk, respectively (Figure 4). No ZnP was detected on uncoated grains (n = 30).

**Figure 4.**
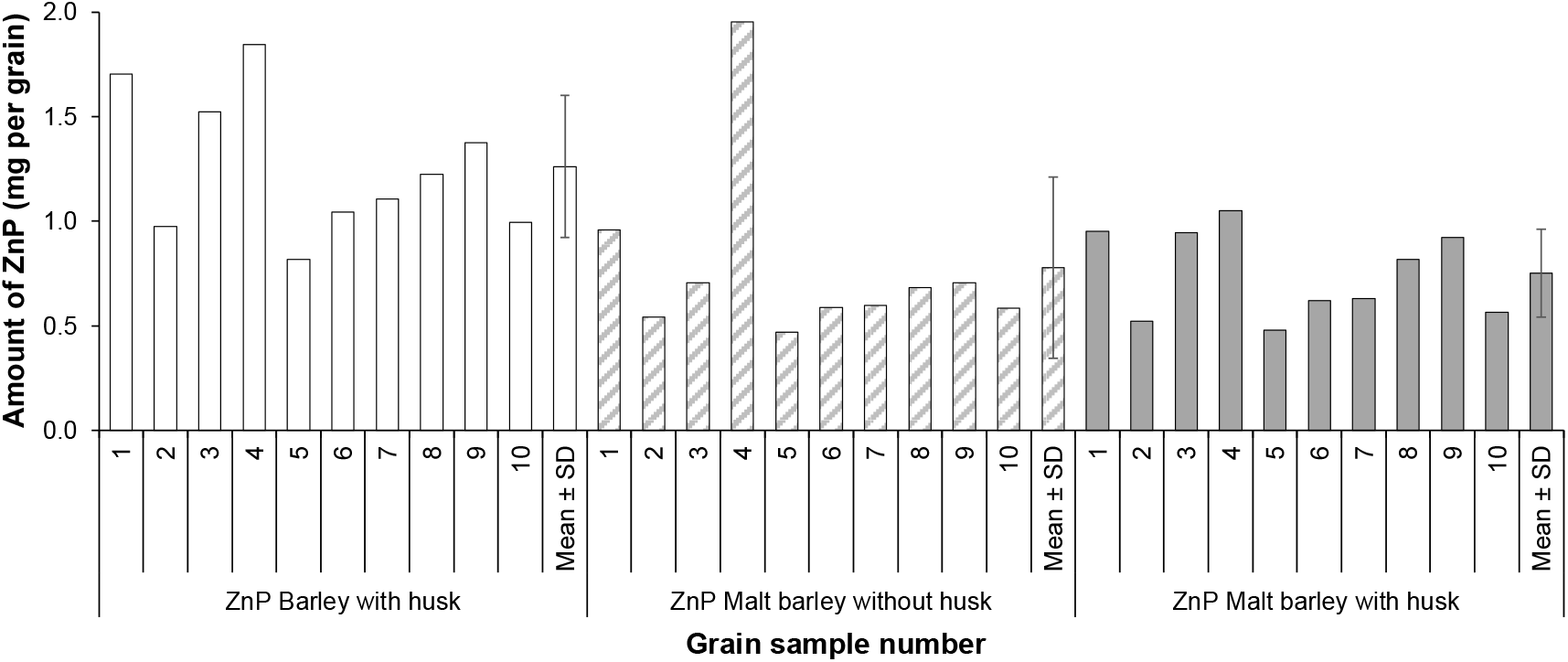
Amount of ZnP on individual grains (mg per grain) (n = 10 grains analysed per grain type), and overall average (Mean ± SD) for each type.

## 4 DISCUSSION

Our laboratory-based study has shown that wild house mice readily switched from a background food to an alternative food and preferred cereals (wheat or barley) over lentils. The preference by house mice for wheat grains is known [23,24,26], but our results indicated that barley was equally favoured. Robards and Saunders [23] included barley in two of 22 pen experiments where different combinations of four choices of grains were offered and found that it performed poorly. Our results showed that it did not matter what single type of background food mice were exposed to, they readily switched to an alternative food on the first night of presentation. Given the neophilic behaviour of mice [30], this is not unexpected. We are not aware of instances where this rapid switching has been reported in the literature for house mice. The background grain type strongly influenced the proportion of alternative grain consumed by mice. If mice had an alternative food type, they avoided lentils. Robards and Saunders [23] found that mice preferred soft wheat varieties over hard varieties. We also observed this in that mice preferred common wheat, a softer grain, to the harder durum wheat variety. Given that there was no difference between wheat and barley grain consumption, barley (malted, with or without husk on) may be a suitable alternative to sterilised wheat as a grain for addition of ZnP in commercial products, but there was no clear advantage over wheat.

This study clearly showed mice consumed toxic ZnP barley grains regardless of the type (barley with husk, malted barley without husk or malted barley with husk). When a less-favoured background grain was available (lentils), the mortality rate of mice was much higher (86%) compared to when a more-favoured background cereal grain, either barley or wheat, was available (mortality of 53% and 47% respectively). This finding suggests that the type of background food was important in determining the choice made by mice when an alternative (toxic bait) grain was made available. Determining an alternative grain for use as a bait in those crops that are more favourable to mice warrants further study. Although there was no apparent difference in mortality rates for the different ZnP-coated barley grains, the type of background food was important in determining the choice made by the mice when a new alternative, albeit toxic, grain was made available.

Our results clearly showed that consumption of a single ZnP-coated grain was not always lethal, and even consumption of up to four ZnP-coated grains did not lead to death for 40% mice. This is despite the coating of ZnP on grains being approximately 1 mg which equates to an LD_90_ dose for mice [31]. However, some coated grains had as little as ∼0.5 mg ZnP/grain while others had up to almost 2 mg ZnP/grain. The results suggest that at the mixing rate of 25 g ZnP/kg grain, on average, most mice would need to consume more than one toxic grain and perhaps more than four toxic grains before receiving a lethal dose. This could explain the low efficacy of ZnP baiting being reported in the field, especially if baiting occurred in the presence of abundant, more-favoured, background food. Further research is required to assess the lethal dose rate of ZnP for Australian house mice as it appears from our findings that the lethal dose rate is likely much higher than 25 g ZnP/kg grain.

In addition, there was a strong and rapid behavioural aversion in mice which did not consume a lethal dose on the first night of exposure to ZnP-coated grains. These mice rapidly switched back to eating their background food, a response which confirms the concern that mice become bait shy if they eat a sub-lethal dose of ZnP [32], but not how rapidly aversion occurs. This parallels the rapid decline in consumption of toxic ZnP bait also observed in common voles, *Microtus arvalis*, after their first night of exposure [17]. The likelihood of any bait shyness compounds the importance of finding the correct bait dosage and indicates the need for strategies to reduce the amount of spilled grain after harvest as noted above.

Most mice died within about 24 h of consumption of ZnP-coated grains, although several animals (14%) died between 39 and 112 h later. This bimodal pattern of mortality reflects the previously reported acute action of ZnP in some mice and the more prolonged effects that reflect organ damage in others (Khan and Schell, https://www.msdvetmanual.com/toxicology/rodenticide-poisoning/zinc-phosphide). Our observations of the clinical signs of toxicity due to ZnP poisoning reflect strongly those described previously by Mason and Littin [32].

Our laboratory experiments included more background food than that required for metabolic maintenance and raises questions about how wild mice forage. For example, in the complex conditions found in farmer’s fields (e.g. growing crops, crop stubble, weeds, other food sources, [33], it is unknown how mice might locate and select the food they consume, including poisoned grains when applied at 1 kg/ha (2-3 grains/m^2^). In zero- and no-till grain cropping systems, spilt grain remaining on the ground immediately post-harvest has been estimated at 20-130 kg/ha (up to ∼390 grains/m^2^), with degradation occurring progressively in the subsequent three to four months to less than 4 kg/ha (up to ∼12 grains/m^2^; Ruscoe et al. unpublished data). Therefore, even if ZnP application rates were higher, it is likely that mice may not encounter ZnP-treated grain amongst existing spilt grain or other abundant food sources in the field. This means that farmers need strategies to improve the effectiveness of ZnP bait against varying background food quantities to prevent high application rates or repeated applications. Understanding these factors and the roles they play in bait uptake, require further research, especially in complex CA systems, and in situations where abundant alternative food exists.

## 5 CONCLUSION

This laboratory-based study has shown that wild house mice will rapidly switch their consumption of one grain type to an alternative on the first night of presentation, and that they prefer cereal grains over lentils. We have also demonstrated that wild house mice will consume different types of barley grains (common barley, malted barley with or without husk) coated with approximately 1 mg ZnP, but the efficacy of this dose is only about 50% when presented as an alternative to a cereal grain compared to in the presence of lentils (87% mortality). Consumption of a sub-lethal dose of ZnP-coated grain also resulted in rapid development of behavioural aversion. We conclude that the currently registered dose rate of ZnP (25 g ZnP/kg wheat; ∼ 1 mg ZnP per grain) in Australia should be re-evaluated to determine what factors may be contributing to variation in efficacy. Further field research is also required to understand the complex association between ZnP dose, and quantity and quality of background food on efficacy of ZnP baits.

## ACKNOWLEDGEMENTS

This work was supported by the Grains Research and Development Corporation (GRDC) through project CSP1804-012RTX, with support from CSIRO Agriculture & Food. We sincerely thank farmers for allowing us to trap mice on their properties for this experiment. We thank Ken Young and Leigh Nelson (GRDC) for ongoing support. All experiments and procedures were approved by the CSIRO Wildlife, Livestock and Laboratory Animal - Animal Ethics Committee and conform to the Australian Code of Practice for the Care and Use of Animals for Scientific Purposes (Approval No 2017-28 and 2018-22).

